# Investigating demic versus cultural diffusion and sex bias in the spread of Austronesian languages in Vietnam

**DOI:** 10.1101/2024.03.19.585662

**Authors:** Dinh Huong Thao, Tran Huu Dinh, Shige Mitsunaga, La Duc Duy, Nguyen Thanh Phuong, Nguyen Phuong Anh, Nguyen Tho Anh, Bui Minh Duc, Huynh Thi Thu Hue, Nguyen Hai Ha, Nguyen Dang Ton, Alexander Hübner, Brigitte Pakendorf, Mark Stoneking, Ituro Inoue, Nguyen Thuy Duong, Nong Van Hai

## Abstract

Austronesian (AN) is the second-largest language family in the world, particularly widespread in Island Southeast Asia (ISEA) and Oceania. In Mainland Southeast Asia (MSEA), groups speaking these languages are concentrated in the highlands of Vietnam. However, our knowledge of the spread of AN-speaking populations in MSEA remains limited; in particular, it is not clear if AN languages were spread by demic or cultural diffusion. In this study, we present and analyze new data consisting of complete mitogenomes from 369 individuals and 847 Y-chromosomal single nucleotide polymorphisms (SNPs) from 170 individuals from all five Vietnamese Austronesian groups (VN-AN) and five neighboring Vietnamese Austroasiatic groups (VN-AA). We found genetic signals consistent with matrilocality in some, but not all, of the VN-AN groups. Population affinity analyses indicated connections between the AN-speaking Giarai and certain Taiwanese AN groups (Rukai, Paiwan, and Bunun). However, overall, there were closer genetic affinities between VN-AN groups and neighboring VN-AA groups, suggesting language shifts. Our study provides insights into the genetic structure of AN-speaking communities in MSEA, characterized by some contact with Taiwan and language shift in neighboring groups, indicating that the expansion of AN speakers in MSEA was a combination of cultural and demic diffusion.

## Introduction

The Austronesian language family (AN), encompassing 1256 languages [1] spoken by approximately 360 million people, stretches from Madagascar to Hainan, Southeast Asia, Taiwan, and Near and Remote Oceania [2]. The Austronesian ancestors are thought to have originated in the Yangtze River Delta 9-6 thousand years ago (kya) [3, 4] and then spread to Taiwan. Nine out of ten AN primary sub-branches are exclusive to Taiwan, while all AN languages outside of Taiwan belong to just a single primary sub-branch (Malayo-Polynesian, consisting of more than 1200 languages), strongly suggesting that Taiwan was the source of the Austronesian expansion [2].

AN groups in Island Southeast Asia (ISEA) have been extensively examined from cultural and biological perspectives, contributing valuable data for elucidating the history of this region [5–17]. However, AN ethnic groups in Mainland Southeast Asia (MSEA) have not yet received the same attention [18–25]. In MSEA, AN groups are found in Vietnam, Thailand, and Cambodia but account for only a small proportion of the population (e.g., about 2.11% of ∼70 million people in Thailand [1]. A crucial question concerning the spread of AN-speaking groups in MSEA is the extent to which this was a process of demic diffusion (i.e., migration of AN groups from elsewhere spreading both their languages and their genes) vs. cultural diffusion (i.e., existing MSEA groups adopting an AN language with little genetic mixing with AN groups from elsewhere). Historical records point to the appearance of AN speakers along the coast of Indochina and the Gulf of Thailand around the 5th century BCE [26]; the close relationship of the AN languages of Vietnam with the Malayic branch of the family [2] points to northwest Borneo as the source of this migration [26]. However, whether the nature of the subsequent diffusion of AN languages was demic or cultural continues to be an open question. Here, we address the question of demic vs. cultural diffusion, and potential sex bias in the spread of AN ancestry, by analyzing mtDNA and Y chromosome variation in AN groups from Vietnam.

Vietnam (VN), with its long coastline, occupies a key geographical position in MSEA and is home to 54 ethnic groups speaking languages classified into five language families: Austroasiastic (AA), Thai-Kadai (TK), Hmong-Mien (HM), Sino Tibetan (ST) and Austronesian (AN). There are five recognized AN groups in Vietnam: Cham, Churu, Ede, Giarai, and Raglay, together accounting for ∼1.32% of the national census size (www.gso.gov.vn; accessed the *General Statistics Office of Vietnam* in July 2023). It is thought that the first AN group to arrive in Vietnam were the ancestors of the Cham on the South Central Coast in 500 BCE [27]. From there, the Cham rapidly extended their territory and established the Champa kingdom [27]. In the process, their languages underwent profound contact-induced changes due to the language shift of the autochthonous populations who were politically subordinate to the Cham [28]. Modern Austronesian communities in Vietnam (VN-AN) mainly occupy the mountainous Central Highlands and the South Central coastline. Their social customs, traditions, and family dynamics are related to the ancient Champa and comparable to their counterparts in ISEA [27]. However, the question of demic vs. cultural diffusion in the spread of AN languages in MSEA has not been investigated. To date, the maternal genetic ancestry of the Vietnamese Cham was described based only on the mtDNA HVS region [20], which provides limited resolution, while complete mtDNA genome sequences are available for two groups of Cham from Cambodia [19]. Recent studies examined both uniparental markers and genome-wide data for two AN groups, Ede and Giarai [18, 21, 23, 24]. These findings were compared with other neighboring populations from different language families but not with other AN ethnicities on the mainland due to data scarcity, hindering attempts to trace AN dispersal and settlement in MSEA.

Here, we present a novel set of complete mtDNA genome sequences of 369 individuals and Y chromosome haplotypes based on genotypes for 847 Y-chromosomal SNPs in 170 individuals, encompassing all five AN ethnic minorities and five neighboring AA groups in Vietnam (Fig 1). This new dataset is analyzed with comparative data from AN groups from elsewhere in ISEA and MSEA, and with non-AN groups from Vietnam, in order to evaluate the relative roles of demic vs. cultural diffusion, and potential sex bias, in the spread of AN languages in Vietnam.

**Fig. 1.**
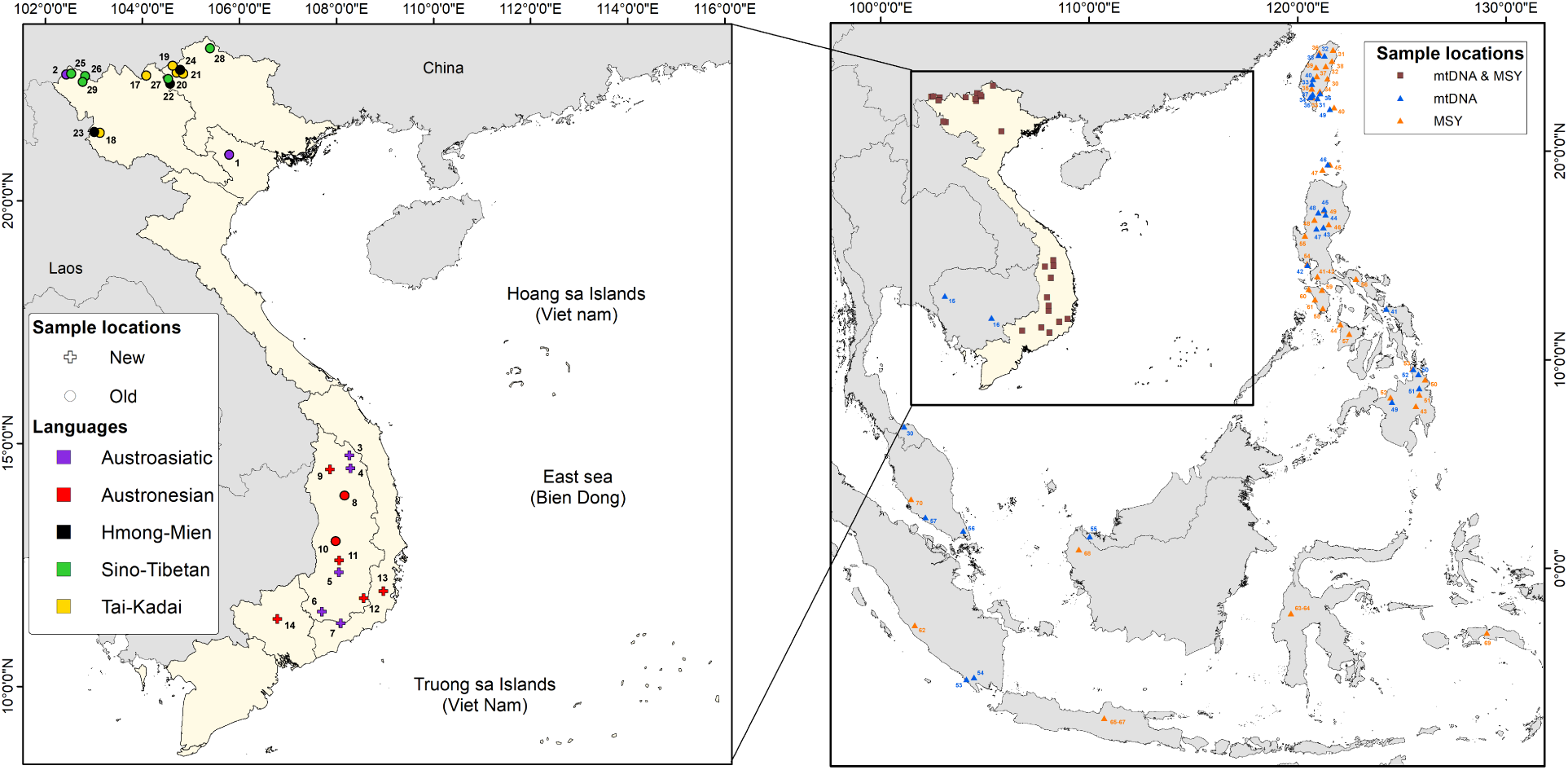
Geographical map displaying sampling locations. Triangles mark the sampling sites of all MSEA and ISEA populations included in the analysis. In the inset, crosses and circles mark the sampling locations of the 10 new and 17 previously published Vietnamese populations, respectively. For the Vietnamese populations, labels are color-coded by language family, as indicated in the legend. All the populations outside of Vietnam included here speak AN languages, and their locations are color-coded by mtDNA and MSY. The comparative populations for mtDNA and MSY analyses are numbered according to supplementary tables S1 and S4, respectively.

## Materials and methods

### Sample information

Blood samples were collected from 369 Vietnamese males belonging to five Austronesian-speaking groups (Churu, Raglay, Ede, Giarai, and Cham) and five Austroasiatic-speaking groups (Coho, Hre, Bana, Mnong, and Ma) from the central highlands of Vietnam (Fig 1, S1 Table). These samples were recruited between December 30^th^, 2019 and November 30^th^, 2022. Since Ede and Giarai individuals were also included in a previous study [18], we here distinguish the two sample sets as Ede-I and Giarai-I (taken from Duong et al. 2018) and Ede-II and Giarai-II (this study). Similarly, in order to distinguish the Cham individuals from Vietnam (this study) from Cham individuals from Cambodia included in Kloss-Brandstätter et al. (2021), we label them Cham-VN and Cham-CB, respectively, further distinguishing the Cambodian samples from Battambang (Cham-CB-Bat) from those from Kampong (Cham-CB-Kam). All participants gave written, informed consent to donate blood, were unrelated, and self-identified to have at least three generations of the same ethnicity. This study received ethical approval from the Institutional Review Board of the Institute of Genome Research, Vietnam Academy of Science and Technology (No: 2-2019/NCHG-HĐĐĐ).

### mtDNA sequencing

Genomic DNAs were extracted with the GeneJET Whole Blood Genomic DNA Purification Mini Kit (ThermoFisher Scientific, USA) following the manufacturer’s protocol. Construction of genomic libraries and capture enrichment for mtDNA were performed as described previously [29]. The libraries were sequenced on the Illumina platform, and the reads underwent quality control and were processed as described previously [30]. Reads were aligned to the Reconstructed Sapiens Reference Sequence (RSRS) [31] using an in-house alignment program, and a multiple sequence alignment was performed using MAFFT [32]. The mtDNA haplogroups were classified by HaploGrep2 [33] with PhyloTree mtDNA tree Build 17 [34]. Haplogroups labeled with an asterisk exclude all downstream subhaplogroups, whereas haplogroup labels without an asterisk include all downstream subhaplogroups. For instance, M71 refers to all haplotypes belonging to haplogroup M71, while R* refers to haplotypes assigned to R, but not assigned to any of the defined subhaplogroups within R. For subsequent analysis, except for haplogroup identification, we excluded positions with missing nucleotide (Ns) and the following sites: poly-C stretch of hypervariable segment 2 (HVS-II; nucleotide positions (np) 303–317); CA-repeat (np 514–523); C-stretch 1 (np 568–573), 12S rRNA (np 956–965), historical site (np 3,107), C-stretch 2 (np 5,895–5,899), 9 bp deletion/insertion (np 8,272–8,289), and poly-C stretch of hypervariable segment 1 (HVS-I; np 16,180–16,195). For comparative analyses, we included 1598 complete mtDNA sequences of 47 populations from Taiwan, ISEA (Philippines, Indonesia, and Malaysia), and MSEA (Vietnam, Cambodia, and South Thailand) [10, 18, 19, 35–39] (S1 Table).

### Y-Chromosomal SNP genotyping and data analysis

A total of 170 samples from the 10 above-mentioned Vietnamese populations were genotyped using the Affymetrix Axiom Genome-Wide Human Origins array. We then extracted a total of 2088 SNPs on the non-recombining region of the Y chromosome (MSY) for this study (S1 Dataset and S2 Dataset); analyses of the autosomal SNP data for these individuals are part of a further study. For subsequent analysis, these SNPs were overlapped with ∼2.3 million bases of the MSY of 600 previously published Vietnamese samples [24], resulting in 2079 SNP positions (S3 Dataset and S4 Dataset). Positions with two or fewer supporting reads were marked as missing. After removing these sites, 847 SNPs remained; the final SNP genotypes and their positions on hg19 are provided in Supplementary Materials (S5 Dataset and S6 Dataset). Genotypes were then used to identify haplogroups by yhaplo [40] using a stopping condition parameter “ancStopThresh” = 10. Haplogroups were determined to the maximum depth possible, given the phylogeny of ISOGG version 11.04 (http://www.isogg.org/) and the SNPs for which we had data. Labels denoted with an asterisk in the text and figures are paragroups that do not include subgroups. Two previously published samples, Kinh09 and Mang304, were excluded from this study as their haplogroups changed to non-informative ancestral haplogroups with the reduced set of SNPs used in this study. For the comparative dataset outside of Vietnam, we calculated the frequencies of haplogroups of 1081 samples from 58 AN-speaking populations from Taiwan and ISEA (Philippines, Indonesia, and Malaysia) [41, 42] based on reported Y-chromosomal SNP datasets (S4 Table). We use the term haplotype throughout this paper to refer to the Y chromosome SNP sequences and not to STR profiles.

### Data analysis

For both uniparental markers, we calculated the number of unique haplotypes for each population with the R function haplotype (package: pegas). Summary genetic statistics of mtDNA and Y chromosome variation, including haplotype diversity (*H*), nucleotide diversity (π), and its variance, were calculated by the functions hap.div and nuc.div of the same package, respectively. To visualize π and *H*, we computed the percentage difference from the mean for each population. The mean number of pairwise differences (MPD) was obtained by averaging over the sum of nucleotide differences for each pair of sequences within a population (R function: dist.dna, package: ape) divided by the total number of pairs. The haplotype sharing within and between populations was estimated as the proportion of pairs of identical sequences shared between populations using an in-house R script. Arlequin version 3.5.2.2 [43] was used to calculate the pairwise genetic distance (ΦST distances) among the populations. For mtDNA markers, we used ΦST distances for computing a nonmetric multidimensional scaling (MDS) analysis (function: isoMDS, package: MASS). The correspondence analysis (CA) was computed based on haplogroup frequencies in R using libraries “vegan” and “ca”.

## Results

### Haplogroup classification of mtDNA and Y-chromosomal SNPs

The entire mitochondrial genomes of 369 individuals were sequenced with an average read depth of 312X (range: 26-3357); 222 distinct sequences (haplotypes) were obtained and assigned to 65 haplogroups by HaploGrep2, all belonging to the two macro-haplogroups M and N (S2 Table). Of the 65 haplogroups, 17 (26.15%) were singletons (S2 Table). Overall, most sequences belonged to the macro-haplogroup M (54.74%), followed by haplogroups B (18.16%) and F1 (13.55%) (S3 Table, S1A Fig). However, there are notable differences in haplogroup frequencies between different MSEA-AN populations, and even between sub-samples from the same ethnolinguistic group, e.g. the Giarai-I and Giarai-II, and also the Cham from Vietnam and those from Cambodia (S2 Fig). The Giarai-I and Ede-I and -II have high frequencies (37-42%) of M71, which is practically absent in other MSEA-AN populations, including the Giarai-II, which were sampled in a location that is about 75 km away from Giarai-I. Similarly, M24b, which is practically absent in other AN populations from MSEA, is found in the Raglay at 35% frequency; haplogroup B, which is found at 17% frequency on average in the other MSEA-AN populations, is found in only very low frequencies in Giarai-I (4%) and Raglay (3%; S1 Table).

For the Y chromosome, a total of 18 polymorphic sites were found; these defined 17 haplotypes belonging to 16 haplogroups (S5 Table). Seven of the 16 haplogroups were shared between at least 2 populations (S6 Table). Of the 16 haplogroups, 5 (31.25%) are singletons. Overall, O1b is the overwhelmingly predominant haplogroup (74.71%), in particular subhaplogroup O1b1a1a (O-M95) (72.35%), followed by O2a (8.82%), R* (4.12%), and R1a (4.12%) (S5 Table, S1B Fig). However, there are some striking differences in haplogroup frequencies among AN populations. For example, O-M95 is very common in VN-AN populations (42-90%), but occurs at somewhat lower frequencies (14-50%) in Indonesian and Malaysian AN populations, lower still (10%) in a Taiwanese AN population, and at very low frequencies (2% and 5%) in two Philippine AN populations (S3 Fig). The Cham from Vietnam have a high frequency (31%) of J (specifically, sub-haplogroup J2a1), which is only present at very low frequencies in Philippine AN (3%) and Indonesian AN (2%) and is absent in Taiwanese and Malaysian populations (S3 Fig, S4 Table).

### Genetic diversity

The nucleotide diversity (π) and haplotype diversity (*H*) for mtDNA sequences and Y-chromosomal SNPs were calculated for 958 individuals (369 newly sequenced here and 598 previously published individuals) and 768 individuals (170 newly genotyped here individuals and 598 previously published), respectively (Fig 2, S7 Table). For mtDNA, the newly genotyped Vietnamese groups had an average haplotype diversity value of 0.9545, ranging from 0.926 (Ede-II and Cham-VN) to 0.991 (Ma). The average nucleotide diversity (π) was 0.00218 and did not vary much among groups; Mnong had the highest value (π = 0.00233), and Ede-II the lowest value (π = 0.00193). For the Y chromosome, the average values were 0.2947 and 0.005489 for haplotype and nucleotide diversities, respectively. Raglay had the highest haplotype diversity (H=0.649), while Cham-VN had the highest nucleotide diversity (π = 0.00988). The most homogenous group was Ma (H=0.00, π = 0.00), but the sample size (n=6) was also the lowest for the Ma. The MPDs were strikingly low in Ma and Churu, less than 1.0, which probably reflects the very high frequencies of haplogroup O1b1a1a (O-M95) found in 6/6 Ma and 18/20 Churu individuals, respectively, indicating homogeneity of the Ma and Churu paternal ancestries. While there is a tendency for the VN-AN groups to have higher Y chromosomal diversity than their neighboring AA groups (Fig 2), as expected for matrilocal populations [44], mtDNA diversity does not seem to be lower in VN-AN groups than in other VN groups.

**Fig 2.**
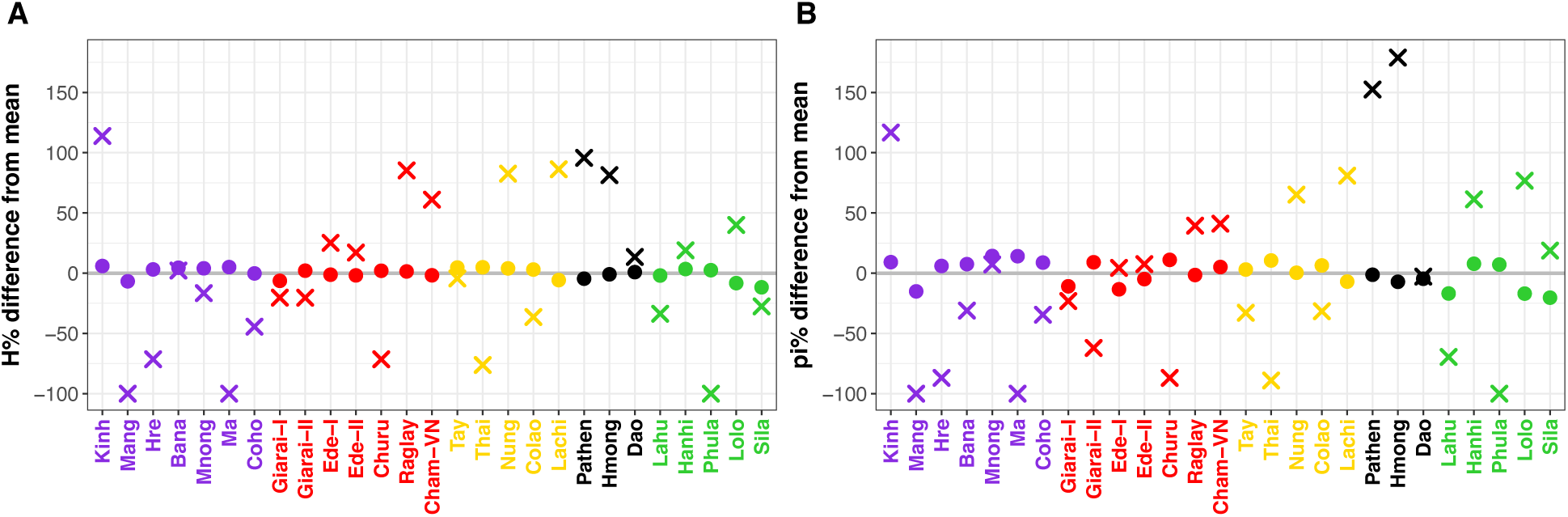
Genetic diversity indices shown as the percent difference from the mean. (A) Haplotype diversity. (B) Nucleotide diversity. Crosses and dots represent MSY and mtDNA data, respectively. Population labels are color-coded by language family, with Austroasiatic in purple, Austronesian in red, Tai-Kadai in yellow, Hmong-Mien in black, and Sino-Tibetan in green. The gray line indicates the mean across populations.

### Population affinity

An mtDNA haplotype sharing matrix was computed for Vietnamese populations and for AN populations from MSEA and ISEA (Fig 3; a similar analysis is not shown for the Y chromosome as there are too few Y chromosome haplotypes to be informative). High values of sharing within groups are an indication of small population size and/or a recent bottleneck, while high values of sharing between populations suggest recent genetic contact or shared ancestry. Among VN-AN groups, within-group sharing is highest in Giarai-I (0.1168) and lowest in Giarai-II (0.0375), comparable to what is observed for non-AN VN groups (0.009 - 0.1206 for VN-AA, 0.0109 - 0.111 for VN-TK, 0.055 - 0.1 for VN-HM, and 0.0246 - 0.1675 for VN-ST; Fig 3). In non-VN-AN populations, higher amounts of intrapopulation sharing are observed in some groups, such as Malaysian Selatar (0.4619), Taiwanese Tao (0.2199), and Malaysian Bidayuh (0.1937). There is some sharing between Vietnamese AN groups and their AA-speaking neighbors, in particular between the Giarai and the Hre and the Bana, between the Ede-I and the Bana and the Ma and the Coho, between the Churu and the Mnong and the Ma and the Coho, and between the Cham-VN and the Ma and the Mnong. However, there is no sharing with other AN populations, with the exception of the two Giarai groups, who share haplotypes with some populations from Taiwan. Furthermore, there is sharing between VN AA-speaking groups Hre and Kinh and Cham from Cambodia.

**Fig 3.**
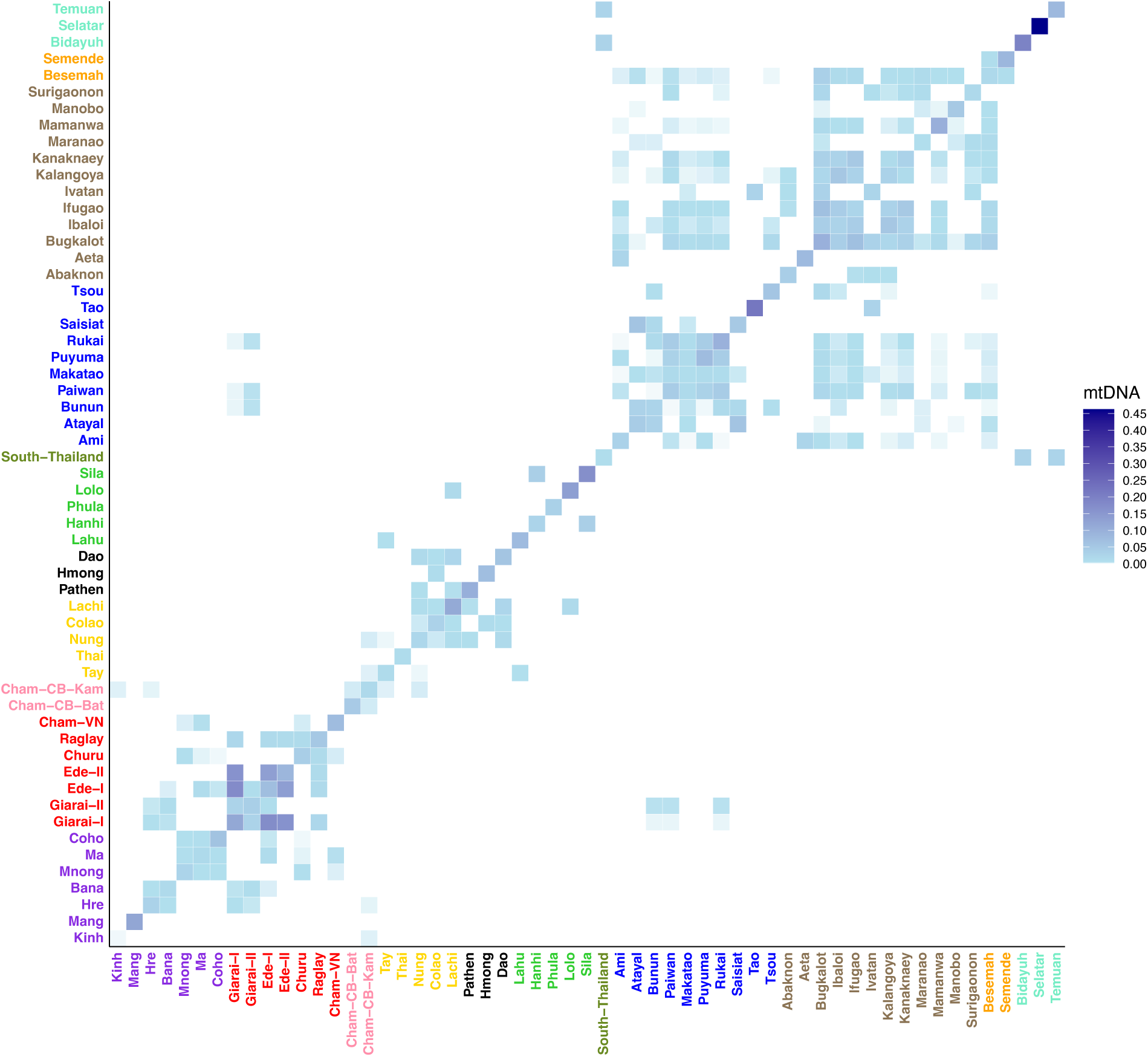
mtDNA haplotype sharing between Vietnamese and non-Vietnamese AN populations. mtDNA shared haplotype frequencies are presented by the blue gradient. Population labels are color-coded by language family with Austroasiatic in purple, Vietnamese Austronesian in red, Thai Austronesian in olive drab, Tai-Kadai in yellow, Hmong-Mien in black, Sino-Tibetan in lime, Cambodian Austronesian in pink, Taiwanese Austronesian in blue, Philippine Austronesian in brown, Indonesian Austronesian in orange, and Malaysian Austronesian in turquoise.

To visualize the relationships among MSEA and ISEA populations, correspondence analysis (CA) was carried out, based on the haplogroup frequencies for both mtDNA and the Y chromosome (Fig 4). For mtDNA, the first dimension differentiated Mang at the top (and to a lesser extent, Pathen and Phula) from Malaysian Austronesian at the bottom (to a lesser extent with southern Thai Austronesian, Indonesian Austronesian, Lolo, and Lachi) (Fig 4A). The second dimension separated most of the Philippine and Taiwanese Austronesian groups at the left from the VN-AN and VN-AA groups at the right. Within this dimension, Cham-VN is pulled towards Taiwanese populations; Cham-CB is grouped with the Southern VN populations, with Cham-CB-Bat being close to the Cham-VN while Cham-CB-Kam is close to the VN-AN Giarai-II and the Churu. All other groups are more towards the middle. On the Y-chromosome CA plot, the first dimension separated most of the Philippine AN groups, including 6 Negrito groups (Mamanwa, Aeta-Bataan, Aeta-Zambles, Agta-Iriga, and Iraya) [41]. The second dimension separated Philippine and Taiwan AN groups at the left from VN-AN and VN-AA groups at the right, similar to the mtDNA plot, except that some VN-TK and VN-ST groups also cluster with the VN-AN and VN-AA groups (Fig 4B). Indonesian AN groups tend to be in the middle in both the mtDNA and Y plots.

**Fig 4.**
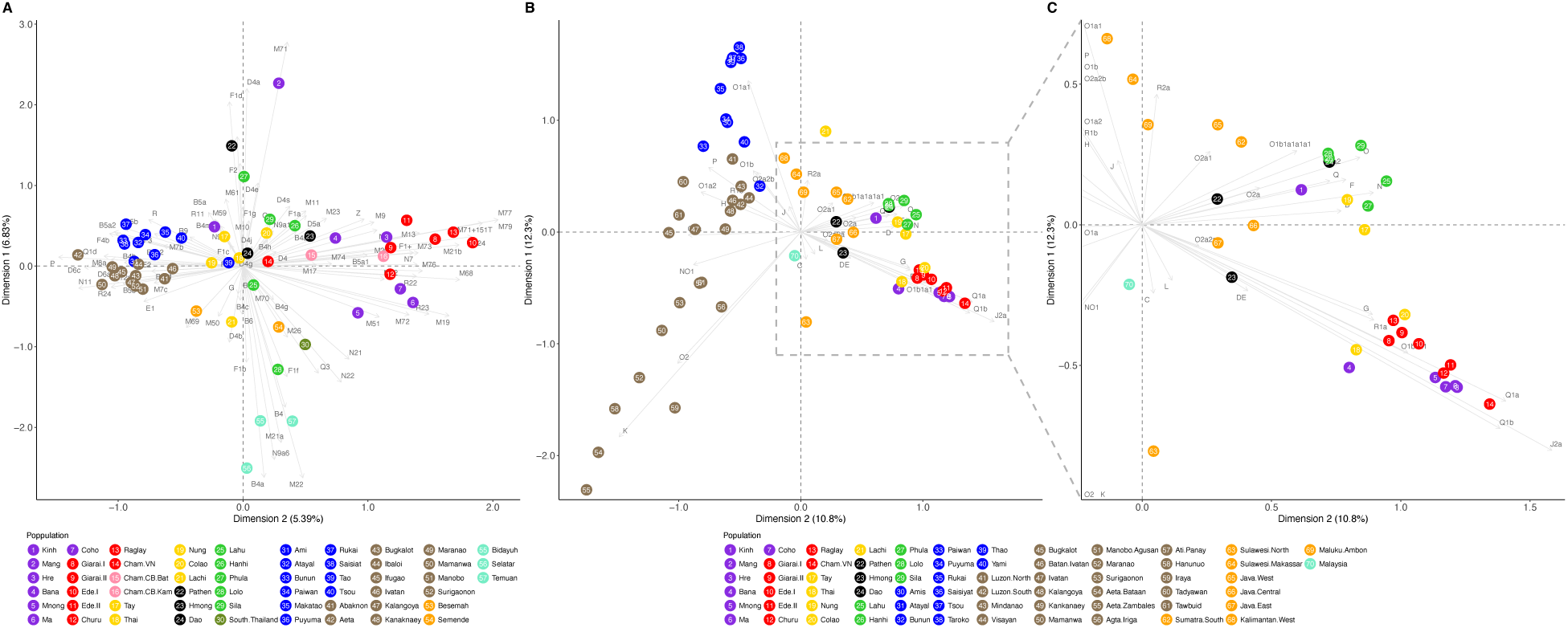
Correspondence analysis plot based on haplogroup frequencies of Vietnamese and non-Vietnamese populations. (A) mtDNA. (B, C) MSY, with the plot on the right (C) zooming in on the region indicated by the dashed rectangle in the full plot (B). Population labels are color-coded by language family with Austroasiatic in purple, Vietnamese Austronesian in red, Tai-Kadai in yellow, Hmong-Mien in black, Sino-Tibetan in lime, Cambodian Austronesian (Cham-CB-Bat and Cham-CB-Kam) in pink, Thai Austronesian in olive drab, Taiwanese Austronesian in blue, Philippine Austronesian in brown, Indonesian Austronesian in orange, and Malaysian Austronesian in turquoise. Haplogroup labels are in gray.

To further investigate genetic differences between populations, pairwise ΦST distances based on mtDNA sequences were generated among Vietnamese populations and AN-speaking populations in other countries (S4 Fig). It was not possible to carry out a similar analysis for the MSY as we only have haplogroup frequencies from the published data, not comparable sequence/SNP data. Among the VN populations, closer relationships (as demonstrated by non-significant ΦST values) are found between the Giarai-II and the AA-speaking Hre, between the Giarai-I and the Ede-I, between the Ede-I and the Ede-II, and between the Cham-VN and the TK-speaking Thai. Among VN and non-VN-AN populations, a non-significant ΦST value is found between the Cham-VN and the Cham-CB-Bat.

An MDS analysis was carried out, using the matrix of pairwise ΦST distances for mtDNA (S5 Fig), to further examine the genetic relationships of the AN groups from Vietnam to other populations from Vietnam and to AN populations from MSEA and ISEA. Because a high stress value was obtained with a two-dimensional analysis, we increased the dimensions to three. The results indicate that the Ede-I and Giarai-I are close to the Taiwanese Atayal and Saisiat in the first dimension, while the Cham-VN and Churu are separate from these and closer to the Cham-CB groups; Ede-II, Giarai-II, and Raglay show affinities with a variety of groups, including AA-speaking Bana and Hre, Taiwanese Ami, Indonesian Besemah, and the Manobo from the Philippines (S5 Fig). In the second dimension, the VN-AN groups are in a central cloud that includes various groups (both AN and non-AN), while the third dimension places the VN-AN groups close to the Cham-CB groups and the VN-AA groups, as well as some other groups from Vietnam.

## Discussion

The five Austronesian-speaking groups in Vietnam (Cham, Raglay, Ede, Giarai, and Churu) share many cultural and social practices [45], yet their genetic relationships with one another, with other Vietnamese groups, and with other Austronesian groups have not been explored in detail. Here, we generated complete mtDNA genome sequences, and determined Y chromosome haplogroup frequencies based on genotyping 847 SNPs, for all five Vietnamese Austronesian-speaking groups, and from five neighboring Austroasiatic-speaking groups. We then compared these to published data from Vietnamese non-AN groups and non-Vietnamese AN groups, in order to assess the role of cultural vs. demic diffusion, and potential sex-biased migration, in the spread of Austronesian languages in Vietnam.

Patterns of genetic differentiation and haplotype sharing between populations provide insights into the question of demic vs. cultural diffusion in the spread of AN languages in Vietnam. With demic diffusion, we would expect to see closer relationships between VN-AN groups and non-VN-AN groups; with cultural diffusion, we would expect to see closer relationships between VN-AN groups and VN-non-AN groups. Moreover, differences between relationships based on mtDNA vs. the MSY would indicate differences in the maternal vs. paternal relationships of VN-AN groups.

The haplogroup frequency plots for both mtDNA and the MSY (S2 and S3 Figs) indicate that VN-AN groups are, overall, more similar to their neighboring VN-AA groups than they are to AN groups from elsewhere. The correspondence analyses (Fig 4) further support this conclusion: for mtDNA, the VN-AN groups overlap with their neighboring VN-AA groups (and are close to Cham-CB groups); while for the MSY, the VN-AN and VN-AA groups are clustered together (and with two VN-TK groups, Thai and Colao) and apart from other groups. The clustering of the Thai and Colao with VN-AN and VN-AA populations reflects the high frequencies of haplogroup O-M95 in these groups (S3 Fig, S4 Table); however, it is unlikely to reflect recent interactions, since it is not evident in the MSY sequence data nor in the genome-wide data available for a subset of these populations [23, 24].

The close genetic similarity between VN-AN groups and neighboring VN-AA groups would suggest a primary role for cultural diffusion in the spread of AN languages in Vietnam, in accordance with linguistic data suggesting language shifts from autochthonous populations beginning soon after the initial AN settlement [28]. Indeed, historical records indicate the AN-speaking Champa maintained close relations with Mon-Khmer local groups, forming both social-economic partnerships and marital alliances [27]. The close genetic relationship between the Cham from Cambodia and the Cham from Vietnam that we see in the mtDNA data mirror their close linguistic relationship: the two groups speak the same language, although their dialects have by now become mutually unintelligible [28]. The split between the two groups is probably recent, since the Cham are thought to have settled in Cambodia only after the fall of the southern Cham capital of Vijaya in 1471 [46]. That this migration occurred after the incorporation of AA-speaking populations into the Cham population is shown by the sharing of mtDNA haplotypes between the Hre and Kinh from Vietnam and the Cham from Cambodia.

A more detailed analysis of mtDNA haplotypes, based on ΦST distances (S4 Fig) and MDS analysis (S5 Fig), provides further support for close relationships between the VN-AN groups and neighboring VN-AA groups, with non-VN-AN groups generally showing more distant relationships. However, a striking result of the haplotype-based analyses is the sharing of mtDNA haplotypes between the Giarai (both groups) and the Bunun, Paiwan, and Rukai of Taiwan (Fig 3). The latter two are southern Taiwanese groups that, in genome-wide data, show closer relationships to AN groups outside Taiwan than do other Taiwanese groups [47]. This would suggest a role for demic diffusion in the introduction of AN languages to Vietnam. However, the direct connection between Taiwan and Vietnam that emerges from the genetic data is not supported by linguistics, which identifies a close relationship of the VN-AN languages with the AN languages of western Indonesia, and specifically northwest Borneo, as mentioned above [2, 26]. This mismatch points to a very complex history of the spread of AN languages to MSEA, which genome-wide studies might help disentangle.

VN-AN groups are characterized by matrilocal residence practice [45, 48]: the husband often relocates from his home village to that of his wife. As such, Y-chromosome diversity is expected to be higher in matrilocal groups than in patrilocal groups, while the opposite is expected for mtDNA diversity [44]. While the 5 VN-AN populations did exhibit generally higher-than-average Y chromosome diversity values, especially compared to the neighboring AA-speaking populations (Fig 2, S7 Table), the mtDNA diversity values were about the same as in other VN groups. Such departures from the strict predictions of the impact of patrilocality vs. matrilocality on patterns of genetic variation are not unexpected, given that many factors impact genetic variation, thereby complicating interpretations of mtDNA and MSY variation [49]. In the case of VN-AN groups, even though marriages are preferred within communities or between groups with similar cultural practices, intermarriages between neighboring groups often do occur [27, 50]. The Vietnamese fundamental residence pattern is either patrilocal or ambilocal [45]; therefore, admixture between neighboring communities might lead to genetic features inconsistent with their cultural kinship systems. This deviation from the expected pattern of genetic variation with matrilocal residence thus provides further support for the substantial amount of contact with local populations – and hence, cultural diffusion – that was involved in the spread of AN languages in Vietnam.

In conclusion, our survey of the mtDNA and MSY relationships of all of the extant VN-AN groups, along with their AA neighbors, illustrates a complex history of migrations as well as cultural diffusion via language shifts and contact, resulting in the spread of AN languages across Vietnam. We see more or less the same picture for both mtDNA and the MSY, suggesting little sex bias in this process (consistent with a primary role for cultural diffusion); however, we caution that our inferences based on the MSY are more limited than for mtDNA, due to the underlying nature of the data. More in-depth studies of MSY variation, as well as genome-wide data, will provide a more holistic picture of the history of AN groups in Vietnam.

## Acknowledgements

We thank all sample donors for contributing to this research. We thank Bui Quang Thanh, Buon Krong Tuyet Nhung, Vo Thi Bich Thuy and Do Hai Quynh for valuable advice and support. We thank the National Institute of Genetics, Japan and the Max Planck Society for research support.

## Ethics Statement

Blood samples were collected from 369 Vietnamese males belonging to five Austronesian-speaking groups (Churu, Raglay, Ede, Giarai, and Cham) and five Austroasiatic-speaking groups (Coho, Hre, Bana, Mnong, and Ma) from the central highlands of Vietnam. All participants were recruited between December 2019 and November 2022. These participants gave written, informed consent to donate blood, were unrelated, and self-identified to have at least three generations of the same ethnicity. This study received ethical approval from the Institutional Review Board of the Institute of Genome Research, Vietnam Academy of Science and Technology (No: 2-2019/NCHG-HĐĐĐ).

## Data Availability Statement

The complete mtDNA sequences generated in the current study are deposited in GenBank (The mtDNA datasetgenerated during the current study are available inGenBank (https://www.ncbi.nlm.nih.gov/genbank/) with accession numbers XXXXXXXX – XXXXXXXX.

## Supporting information

**S1 Dataset. Chromosomal positions on hg19 for 2088 SNPs**

**S2 Dataset. Genotypes of 2088 SNPs for 170 newly genotyped individuals S3 Dataset. Chromosomal positions on hg19 for 2079 SNPs**

**S4 Dataset. Genotypes of 2079 SNPs for 768 individuals (170 newly genotyped here individuals and 598 previously published)**

**S5 Dataset. Chromosomal positions on hg19 for 847 SNPs**

**S6 Dataset. Genotypes of 847 SNPs for 768 individuals (170 newly genotyped here individuals and 598 previously published)**

