## Supplementary figures and images for "Investigating demic versus cultural diffusion and sex bias in the spread of Austronesian languages in Vietnam"

### S1 Fig

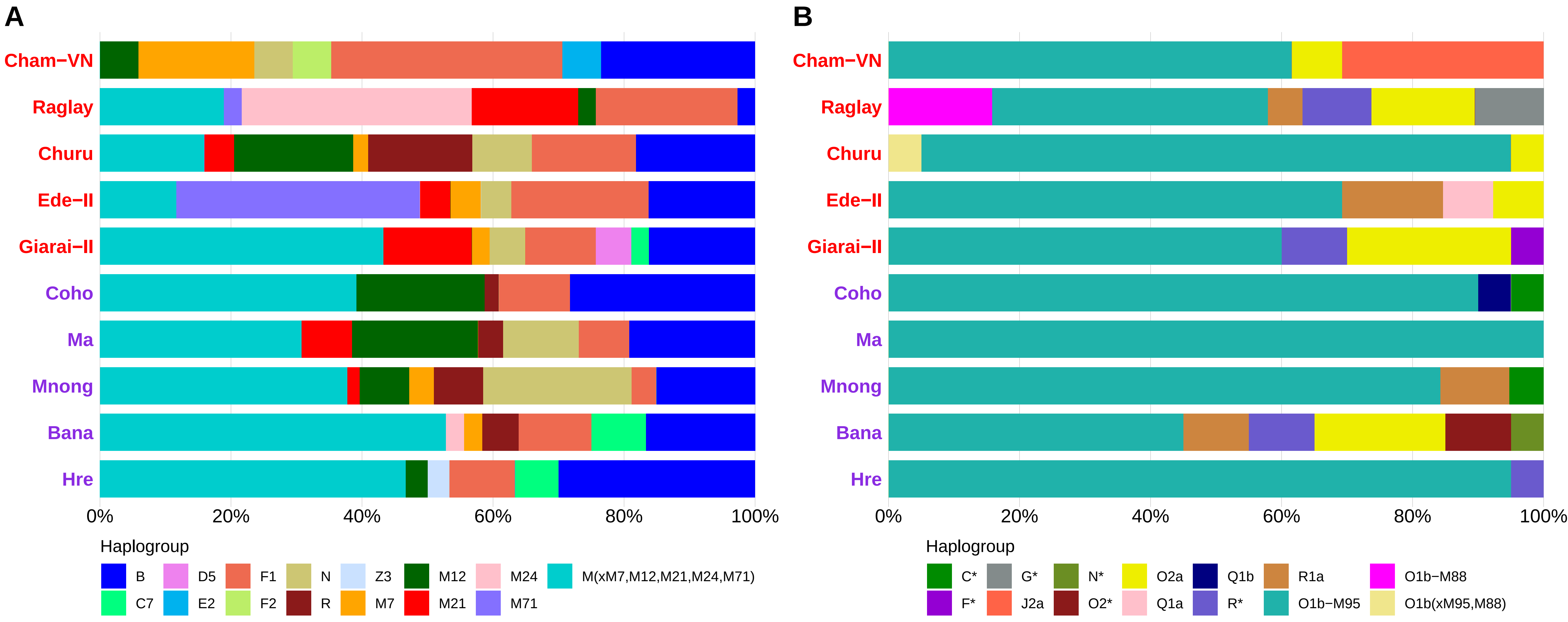

### S2 Fig

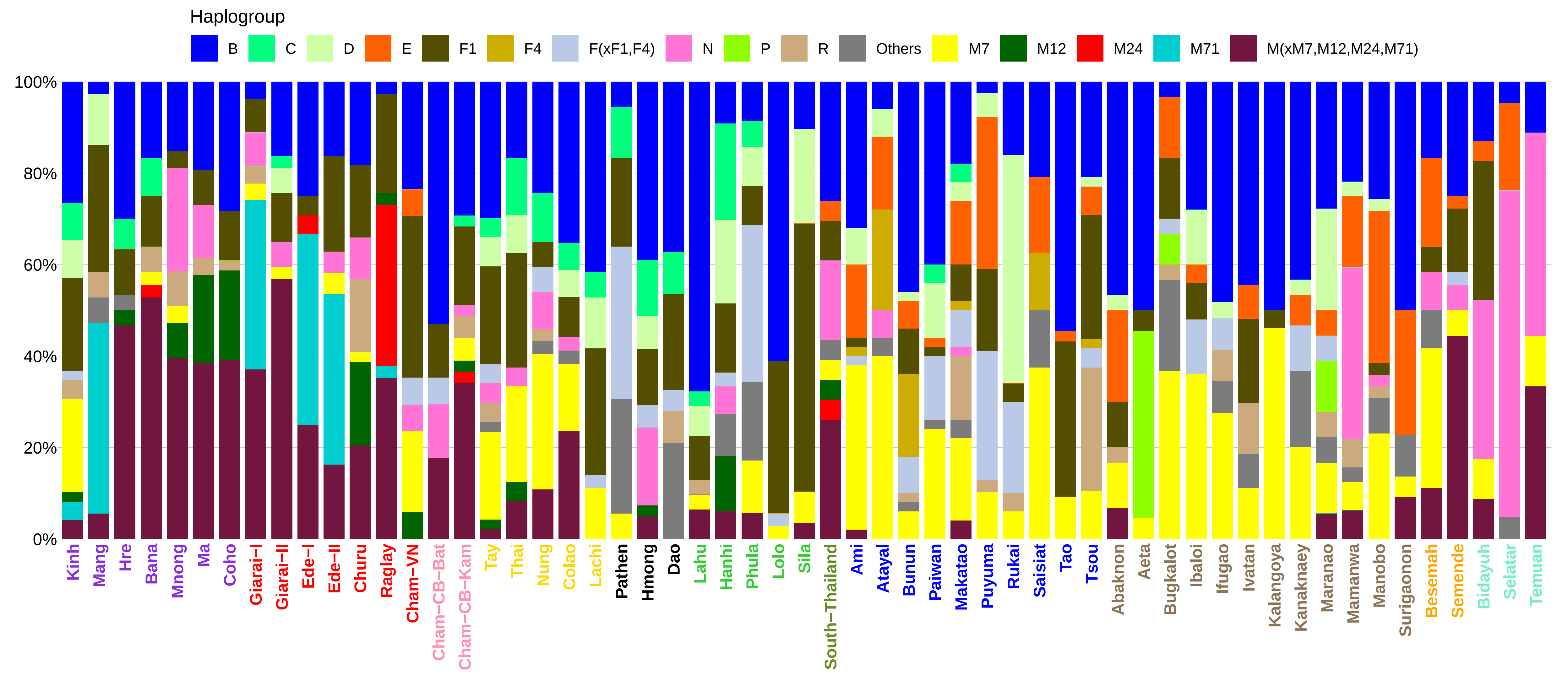

### S3 Fig

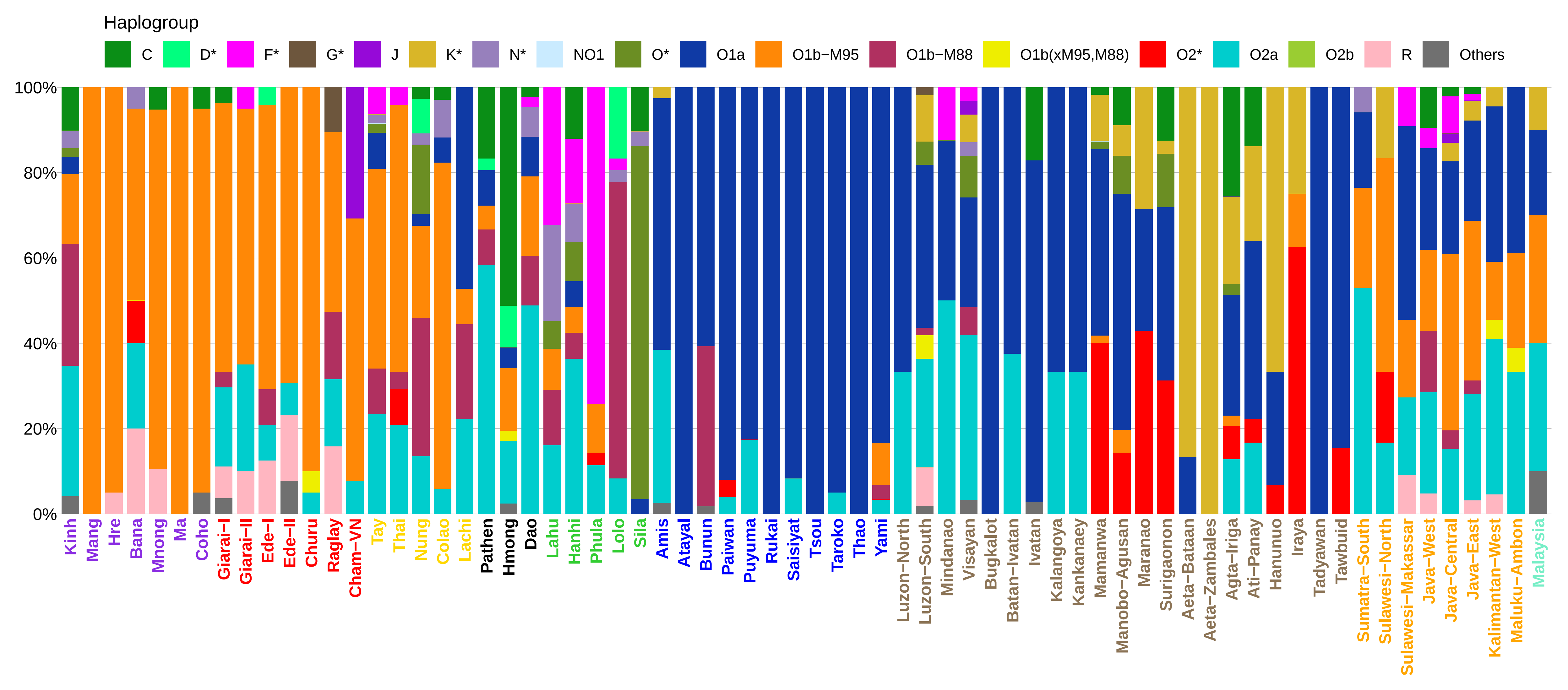

### S4 Fig

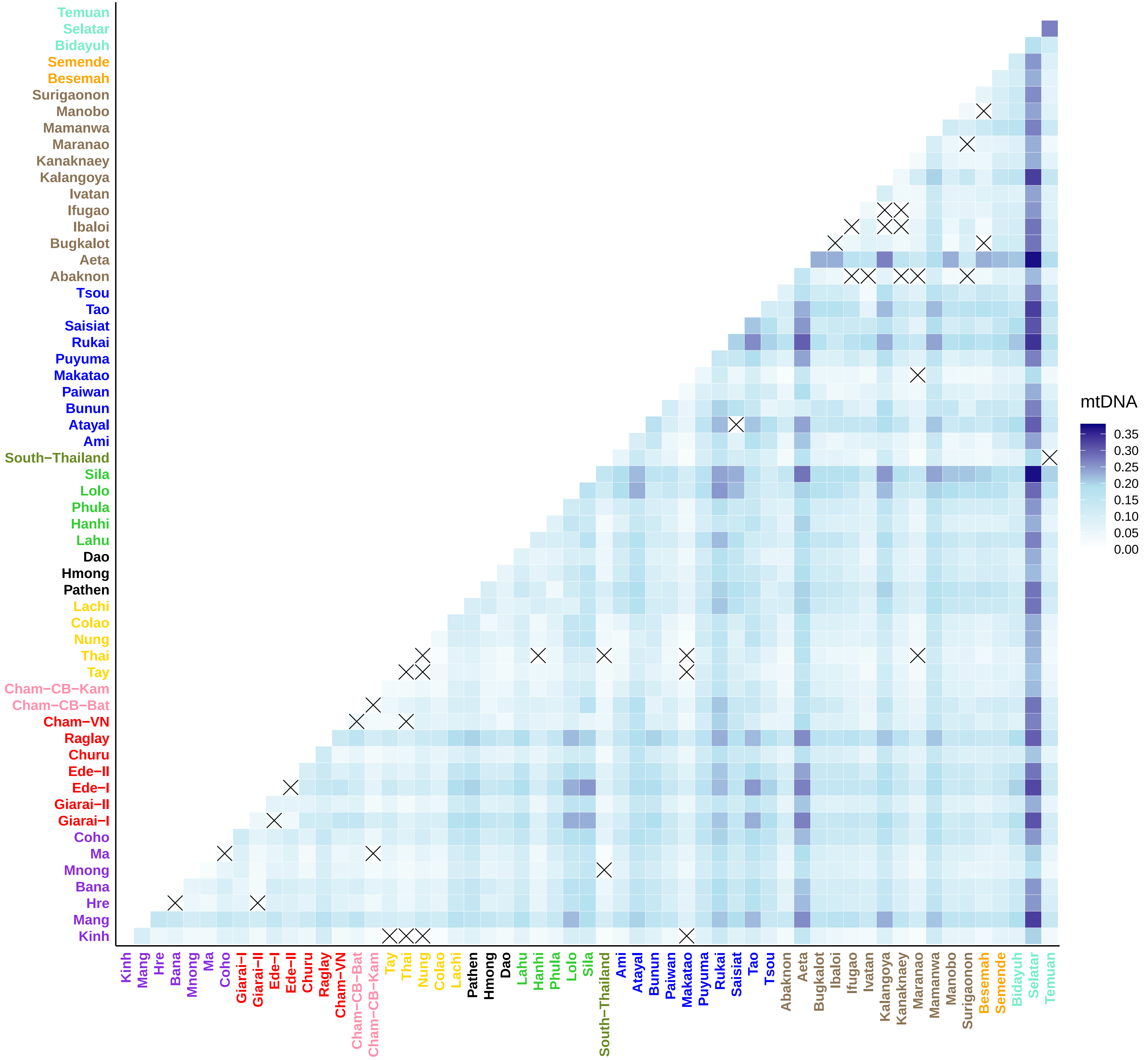

### S5 Fig

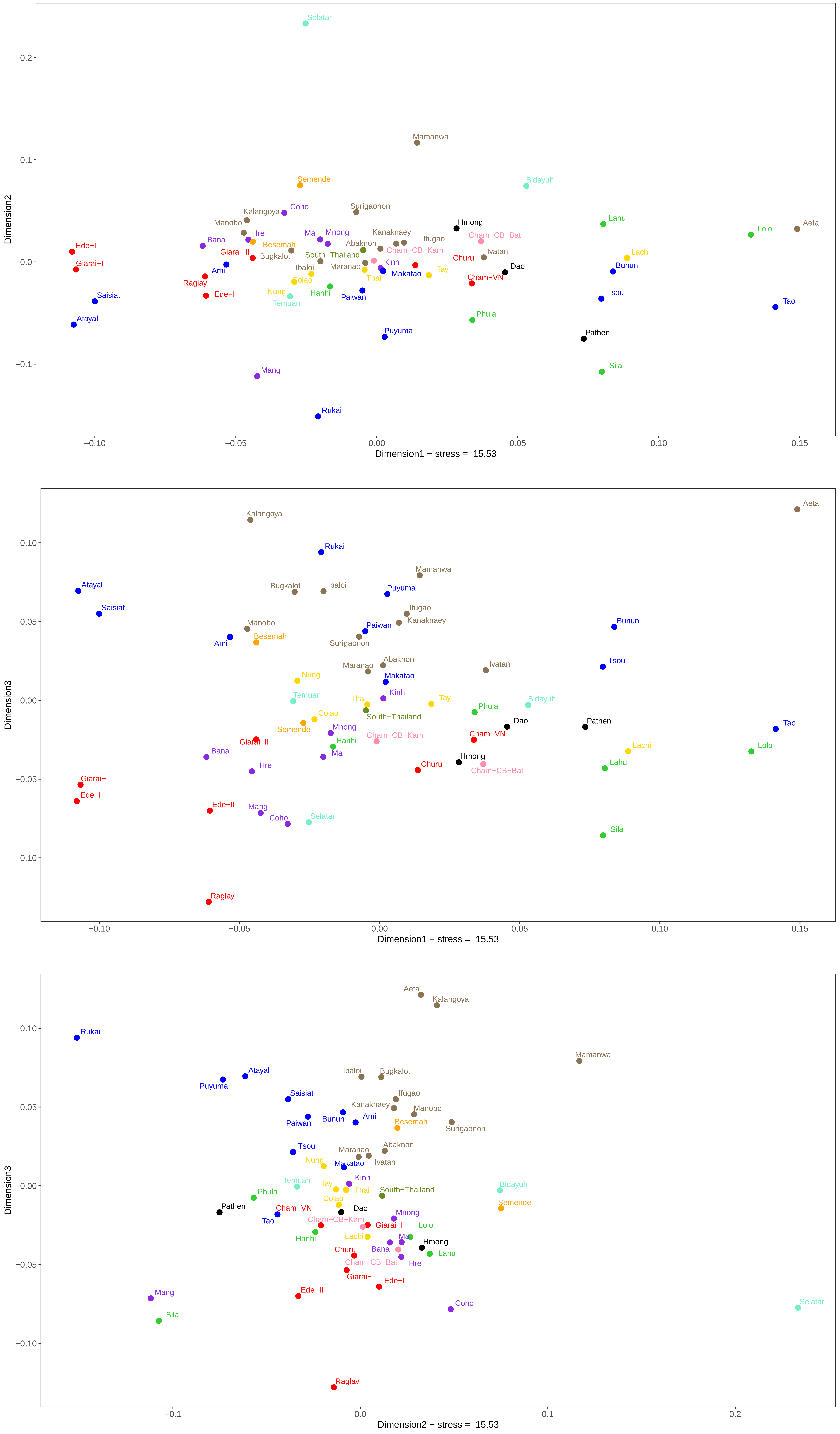
